# Structural bases for blockade and activation of BK channels by Ba^2+^ ions

**DOI:** 10.1101/2022.06.10.495656

**Authors:** Shubhra Srivastava, Pablo Miranda, Natalia de Val, Teresa Giraldez, Jianghai Zhu, Raul Cachau, Miguel Holmgren

## Abstract

Ba^2+^ increases open probability and blocks the permeation pathway of BK channels. Here we discuss Cryo-EM structures of BK channels with Ba^2+^. They reveal that Ba^2+^ occupies site S3 in the selectivity filter, leading to blockade. Ba^2+^ are also detected at all high affinity Ca^2+^ binding sites of the gating ring. However, the architectures of this region in these structures represent intermediate transitions between the extreme close and open configurations.

---

The voltage-dependent and Ca^2+^-activated potassium (BK) channels are unique potassium channels given their large single-channel conductance, which has allowed exquisite characterization of their biophysical properties. An α-subunit of a BK channel consists of a membrane core followed by an intracellular region containing two high-affinity Ca^2+^ binding sites^1–3^. As a tetramer, BK channels form a large intracellular structure with eight high-affinity Ca^2+^ sites^4,5^, called the gating ring^6^. Each Ca^2+^ site is embedded in a regulator of potassium conductance (RCK) domain^7^. Functionally^1–3,8,9^ and structurally^4,5,10^, Ca^2+^ ions interact with BK channels’ RCK1 and RCK2 domains.

Historically, Ba^2+^ was first discovered as a potent blocker of BK channels^11–13^. Recently^14,15^, it has also been shown that Ba^2+^ activate BK channels by interacting with the Ca^2+^-bowl site (RCK2). A unified model integrating these functional properties has not been obtained. Here we discuss two cryoEM structures of *Aplysia* Slo1 BK channels with Ba^2+^ incorporated into nanodiscs. As shown previously^5^, *Aplysia* Slo1 channels display all electrophysiological properties common to BK. Similarly, *Aplysia* Slo1 channels also display blockade and activation by Ba^2+^ (Fig. 1). Fig. 1a shows normalized current levels before and after (arrow) four different Ba^2+^ concentration exposures to the same patch. With 100 μM Ba^2+^, fast blockade of the permeation pathway is observed (open circles). However, exposing the channels to lower Ba^2+^ concentrations, the blockade is slower, unveiling activation by Ba^2+^ that preceded block (black, cyan, and red circles for 10, 5, 1 μM Ba^2+^, respectively). Fig. 1b shows representative current traces for these conditions. As shown previously^15^, activation by Ba^2+^ of *Aplysia* Slo1 channels is weaker than activation by Ca^2+^ (Fig. 1c).

**Fig. 1.**
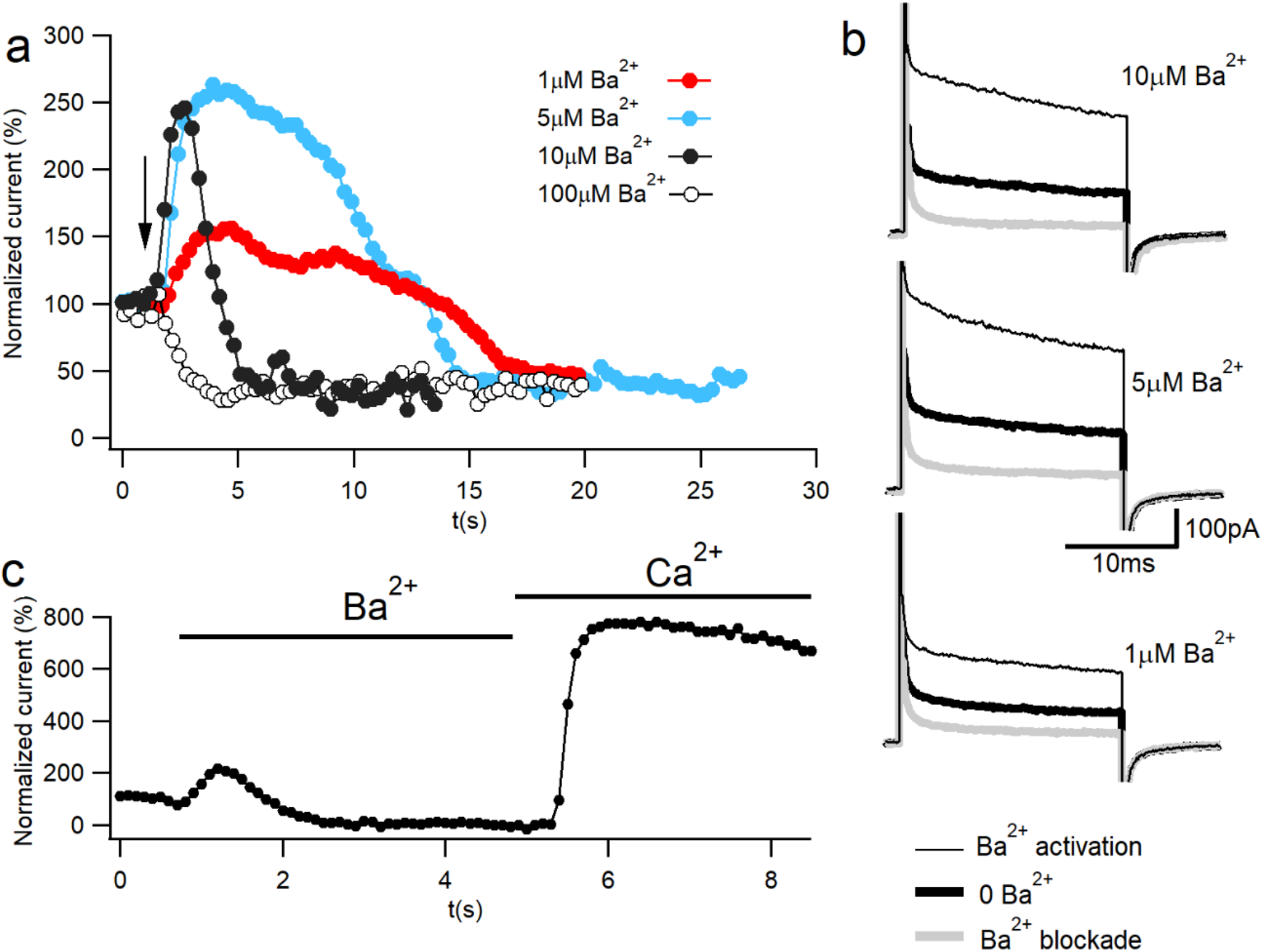
Ba^2+^ activation and blockade of BK current. (**a**) Time course of normalized K^+^ currents in response to 1 μM (red), 5 μM (cyan), 10 μM (full black circle) or 100 μM (open circle) Ba^2+^ applied to an inside-out patch containing *Aplysia* BK channels. Dots represent the average current of the last 10 ms pulse to 120 mV from a holding potential of −70 mV every 300 ms. Arrow represents the moment of the Ba^2+^ addition. (**b**) Representative current traces in 10 μM Ba^2+^ (top panel), 5 μM Ba^2+^ (middle panel) and 1 μM Ba^2+^ (bottom panel). The current in the absence of Ba^2+^ (bold black trace) increases during Ba^2+^ activation (thin black trace) and reduces after Ba^2+^ blockade (gray trace). (**c**) Time course of normalized current showing Ca^2+^ activation after Ba^2+^ activation and blockade. Horizontal bars indicate divalent species applications. Ba^2+^ blockade is reversible and Ca^2+^ activation is 5 times bigger than Ba^2+^ activation.

Previous structures of *Aplysia* Slo1 have been reported in extreme conditions, i.e. in the absence of Ca^2+^ and presence of EDTA (close state; PDBID:5TJI) and in the presence of high concentrations of Ca^2+^ and Mg^2+^ (open state; PDBID:5TJ6)^4,5^. We will refer to these structures as the EDTA and Ca^2+^/Mg^2+^ structures, respectively. Our first structure (PDBID: 7RJT) is in the presence of 10 mM Ba^2+^, while the second (PDBID: 7RK6) was obtained from *Aplysia* Slo1 channel protein purified in the presence of 40 mM Ba^2+^, but after nanodisc incorporation it was run through the size exclusion chromatography column preequilibrated with a buffer lacking divalent ions and chelators. We will refer to these two structures as the 10 mM Ba^2+^ and the low-divalent-ions structures, respectively. The low-divalent-ion structure may help us identify the initial divalent binding geometry at very low ion concentration.

Extended Data Figs. 1 and 2 show views of the overall structure in the presence of 10 mM Ba^2+^ and low-divalent-ions form, respectively. In both structures, eight sites with strong map densities within the gating rings could be attributed to Ba^2+^ ions. We did not observe any electronic density at the low-affinity Mg^2+^ binding site, indicating that Ba^2+^ cannot substitute Mg^2+^ at this site. Only the 10 mM Ba^2+^ structure showed a ninth Ba^2+^ map density in the selectivity filter, at the traditional K^+^ site S3 (Extended Data Fig. 1e). In this structure, map densities attributable to K^+^ ions were observed at S4 and S2 sites, and an additional at the intracellular entrance of the filter. Remarkably, in BK channels, the presence of internal and external K^+^ lock-in sites was predicted functionally more than 30 years ago^11,12^.

**Fig. 2.**
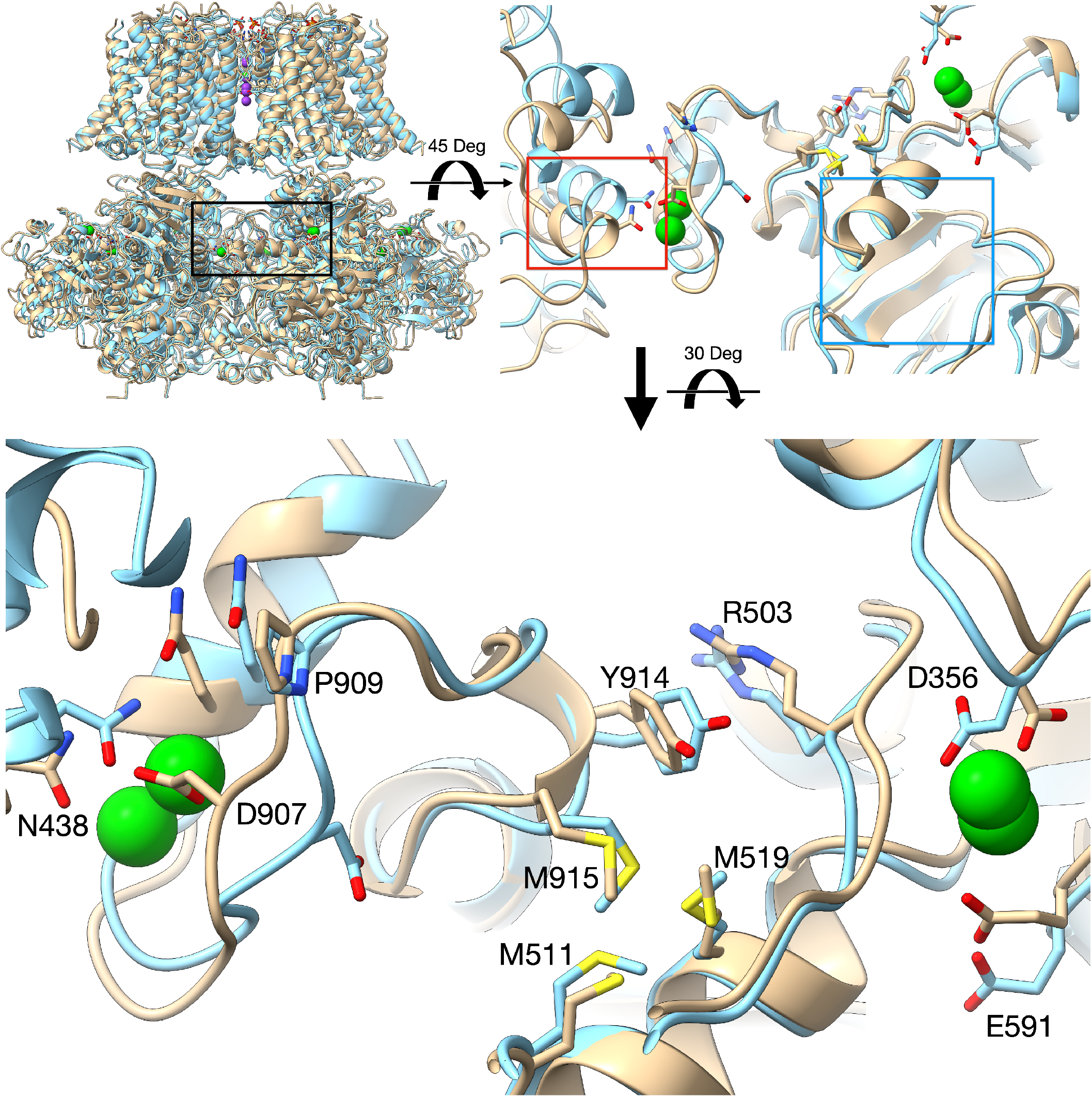
Overview of the 10 mM Ba^+2^ (brown) and low-divalent-ions (light blue) structures (top left) with RCK1/RCK2 region highlighted (black box) and zoomed in (top right). The RCK1/RCK2 of the 10 mM Ba^+2^ (brown) and low-divalent-ions (light blue) structures were iteratively aligned to minimize <RMSD(N)>/N^k^) with N the number of atoms in the region optimizing the RMSD (optimized k=0.47). Notice the near perfect overlap of the beta-sheet structures immediately beneath the RCK1/RCK2 (blue rectangle). The shift of the αD helix from a neighboring subunit is indicated by the red box. The bottom section shows the changes in ion coordination between the two structures.

In the Ca^2+^/Mg^2+^ structure, divalent ions introduce a new bend of the S6 helices at Gly302^4,5^. Since this residue is near the selectivity filter, this conformational change is likely the most consequential movement linked to the opening of the permeation gate in the *Aplysia* Slo1 channel. However, we do not observe this bend in our structures. Interestingly, even though the structures did not show the bend at Gly302, they do reflect intermediate expansions (as measured from the distance between Cα atoms of the two Lys320 across the pore axis) from the limits set by the EDTA (38.8 Å) and the Ca^2+^/Mg^2+^ (52.3 Å) structures^4,5^, showing values of 40.7 Å and 44.7 Å for the low-divalent-ions and the 10 mM Ba^2+^ structures, respectively (Extended Data Figs. 1c, 2c and 3b).

As previously^4,5^, in our two structures, the RCK1 N-lobe showed much larger motions than the RCK1 C-lobe. The helix αB appears to shift towards its C-terminal end in the 10 mM Ba^2+^ structure. This movement is driven by the displacement of the intracellular domain relative to the trans-membrane domain (Extended Data Fig. 3a), with the helix αB in close proximity to the interphase between the two domains. As a result of this movement, one of the Mg^2+^ coordinating residues, E388, moves towards the position found in the Ca^2+^/ Mg^2+^ structure^4,5^ (Extended Data Fig. 3c). These observations help explain previous functional data showing that the Mg^2+^ affinity is higher in open Slo1 channels than in closed ones^16^ and that the ability of Mg^2+^ to enhance channel activation is by affecting the close→open transition^17^. Additionally, Asp421 (N-terminus of helix αD) swings by 5 Å from its position in the EDTA structure to the 10 mM Ba^2+^ structure, about halfway from the final position in the Ca^2+^/Mg^2+^ structure^4,5^, providing further support that our structures represent intermediate transitions between the close and open states. Even though the RCK2 is less flexible than RCK1, the small changes observed in response to Ba^2+^ are comparable to those observed in response to Ca^2+^^4,5^. A particular flexible region is the Ca^2+^ bowl, which we will discuss later.

Most residues interacting with Ca^2+^ at the RCK1 and RCK2 sites in the Ca^2+^/Mg^2+^ structure^5^ interact with Ba^2+^ (Fig. 2). By comparative analysis with previous structures, the positions of the side chains coordinating the Ba^2+^ ions in our structures also reflect successive intermediates between the position in the EDTA and Ca^2+^/Mg^2+^ structures^4,5^. In the Ca^2+^-bowl, the side chain of Asp907 is not facing the binding site pocket even with the (partial) presence of Ba2+ in the binding pocket in the low-divalent-ions structure (Fig. 2), as in the EDTA structure^4^. The small linker between helices 310 and αR, containing Asp907 appears to be quite flexible. In the 10 mM Ba^2+^ structure, the Asp907 side chain faces the Ca^2+^ bowl binding pocket (Fig. 2), as in the Ca^2+^/Mg^2+^ structure^5^. As mentioned earlier^4,5^, this movement of Asp907 in response to Ba^2+^ triggers an intricate series of concerted changes suggesting coordinated dynamics of the two binding sites within the same subunit (Supplementary Video). Key residues involved in this flexible corridor are Pro909, Tyr914, Arg503, the Methionine cluster involving Met511, Met 519 and Met915, and an Asn438 from a neighboring subunit. Some of these residues have been previously identified as key players in functional studies^9^,^18–20^. Interestingly, these rearrangements are facilitated by the rotation of the methionine cluster which acts as a buffer between the dynamics of the two binding sites (Supplementary Video). Beneath the flexible corridor, there is a rigid beta strand structure impeding changes in the direction perpendicular to the membrane (Fig. 2, blue square). Noticeably, the general architecture of this flexible corridor is maintained in the hBK channels^10^. Since the “face-in ↔ face-out” dynamics of Asp907 are not strictly dependent on the presence/absence of Ba^2+^, we hypothesize that they would provide the basis for Ba^2+^ activation through the Ca^2+^-bowl with an ~fivefold reduction in affinity as compared to Ca^2+^(15).

More work is needed to explain the structural bases for the selective nature of Ba^2+^ activation. Clearly it is not the absence of Ba^2+^ at the RCK1 site. We can only speculate that the contribution of Asp356 to the coordination of a divalent ion at RCK1 site^4,5,10^ is crucial to trigger the conformational changes responsible for the activation of the gating ring through the RCK1 domain. This residue shows a rearrangement between our two structures, possibly compensatory of the Arg503 displacement and partial disengagement from the Ba^2+^ coordination sphere at low ion concentration (Fig. 2 & Extended Data Fig. 3d). Yet, this movement is not as large as those observed previously^4,5^, and the side chain of Asp356 did not reach distances from the Ba^2+^ consistent with ion coordination.

## Supporting information

Data Quality

## Notes

### Competing Interest Statement

The authors have declared no competing interest.

### Summary of Updates

Ba2+ increases open probability and blocks the permeation pathway of BK channels. Here we discuss Cryo-EM structures of BK channels with Ba2+. They reveal that Ba2+ occupies site S3 in the selectivity filter, leading to blockade. Ba2+ are also detected at all high affinity Ca2+ binding sites of the gating ring. However, the architectures of this region in these structures represent intermediate transitions between the extreme close and open configurations.

## References

1. Bao, L., Rapin, A.M., Holmstrand, E.C. & Cox, D.H. Elimination of the BKCa channel’s high-affinity Ca^2+^ sensitivity. J Gen Physiol 120, 173–89 (2002).

2. Xia, X.M., Zeng, X. & Lingle, C.J. Multiple regulatory sites in large-conductance calcium-activated potassium channels. Nature 418, 880–4 (2002).

3. Zeng, X.H., Xia, X.M. & Lingle, C.J. Divalent cation sensitivity of BK channel activation supports the existence of three distinct binding sites. J Gen Physiol 125, 273–86 (2005).

4. Hite, R.K., Tao, X. & MacKinnon, R. Structural basis for gating the high-conductance Ca^2+^-activated K^+^ channel. Nature 541, 52–57 (2017).

5. Tao, X., Hite, R.K. & MacKinnon, R. Cryo-EM structure of the open high-conductance Ca^2+^-activated K^+^ channel. Nature 541, 46–51 (2017).

6. Jiang, Y. et al. Crystal structure and mechanism of a calcium-gated potassium channel. Nature 417, 515–22 (2002).

7. Jiang, Y., Pico, A., Cadene, M., Chait, B.T. & MacKinnon, R. Structure of the RCK domain from the E. coli K^+^ channel and demonstration of its presence in the human BK channel. Neuron 29, 593–601 (2001).

8. Bao, L., Kaldany, C., Holmstrand, E.C. & Cox, D.H. Mapping the BK_Ca_ channel’s “Ca^2+^ bowl”: side-chains essential for Ca^2+^ sensing. J Gen Physiol 123, 475–89 (2004).

9. Zhang, G. et al. Ion sensing in the RCK1 domain of BK channels. Proc Natl Acad Sci USA 107, 18700–5 (2010).

10. Tao, X. & MacKinnon, R. Molecular structures of the human Slo1 K^+^ channel in complex with beta4. Elife 8(2019).

11. Neyton, J. & Miller, C. Discrete Ba^2+^ block as a probe of ion occupancy and pore structure in the high-conductance Ca^2+^-activated K^+^ channel. J Gen Physiol 92, 569–86 (1988).

12. Neyton, J. & Miller, C. Potassium blocks barium permeation through a calcium-activated potassium channel. J Gen Physiol 92, 549–67 (1988).

13. Vergara, C. & Latorre, R. Kinetics of Ca^2+^-activated K^+^ channels from rabbit muscle incorporated into planar bilayers. Evidence for a Ca^2+^ and Ba^2+^ blockade. J Gen Physiol 82, 543–68 (1983).

14. Miranda, P., Giraldez, T. & Holmgren, M. Interactions of divalent cations with calcium binding sites of BK channels reveal independent motions within the gating ring. Proc Natl Acad Sci U S A 113, 14055–14060 (2016).

15. Zhou, Y., Zeng, X.H. & Lingle, C.J. Barium ions selectively activate BK channels via the Ca^2+^-bowl site. Proc Natl Acad Sci U S A 109, 11413–8 (2012).

16. Shi, J. & Cui, J. Intracellular Mg^2+^ enhances the function of BK-type Ca^2+^-activated K^+^ channels. J Gen Physiol 118, 589–606 (2001).

17. Zhang, X., Solaro, C.R. & Lingle, C.J. Allosteric regulation of BK channel gating by Ca^2+^ and Mg^2+^ through a nonselective, low affinity divalent cation site. J Gen Physiol 118, 607–36 (2001).

18. Bukiya, A.N. et al. An alcohol-sensing site in the calcium- and voltage-gated, large conductance potassium (BK) channel. Proc Natl Acad Sci U S A 111, 9313–8 (2014).

19. Li, Q. et al. Molecular determinants of Ca^2+^ sensitivity at the intersubunit interface of the BK channel gating ring. Sci Rep 8, 509 (2018).

20. Shi, J. et al. Mechanism of magnesium activation of calcium-activated potassium channels. Nature 418, 876–80 (2002).

