## Extended Data for "Structural bases for blockade and activation of BK channels by Ba^2+^ ions"

### Methods

#### Expression and purification

The gene fragment encoding the full-length Slo1 from *Aplysia californica* was expressed as a fusion construct with a C-terminal GFP/rho-1D4 tag cleavable by precision protease in Sf9 cells[1,2]. The baculovirus was generated in Sf9 cells following the standard Bac-to-Bac system protocol. In brief, transfection of sub-confluent cells was carried out with recombinant Slo1 bacmid DNA using Cellfectin II transfection reagent according to manufacturers' instructions. Cell culture supernatants were collected after four days to obtain P1 recombinant baculovirus. Sf9 cells were infected with P1 virus stock to produce P2 virus, then used for subsequent infection to produce P3 viruses. Finally, the P3 virus was used for recombinant protein production.

Sf9 cells were cultured in SFM III Expression Medium for large-scale protein expression. At a cell density of $2 \times 10^6$ cells per ml, infection was performed with recombinant baculovirus and incubated at 27 °C with agitation. Forty-eight hours after infection, cells were harvested by centrifugation. All purification procedures were carried out at 4 °C. The cell pellet was re-suspended and lysed by osmotic shock in a hypotonic buffer consisting of 10 mM Tris-HCl pH 8.0, 3 mM dithiothreitol (DTT), 1 mM EDTA, and protease inhibitors pepstatin A (0.1 μg/ml), aprotinin (1 μg/ml), leupeptin (1 μg/ml), soy trypsin inhibitor (0.1 μg/ml), phenylmethylsulphonyl fluoride (0.2 mM), benzamidine (1 mM) and 4-(2-aminoethyl) benzenesulfonyl fluoride hydrochloride (0.1 mg/ml). After lysis, cell membranes were collected by centrifugation at 30000 g for 30 min. The pellet was homogenized in a buffer containing 20 mM Tris-HCl pH 8.0, 320 mM KCl, 15 mM $BaCl_2$,

5 mM EGTA, and all protease inhibitors added previously. n-Dodecyl-β-D Maltopyranoside (DDM) and Cholesterol Hemi Succinate (CHS) were added to the cell membrane suspension to achieve a 1% and 0.2% (w/v), respectively. Then, the suspension was stirred gently for 1 hr, followed by ultracentrifugation at 45000 g for 30 min. Clarified supernatant was added to a GFP nanobody-conjugated affinity resin pre-equilibrated with buffer composed of 20 mM Tris-HCl pH 8.0, 320 mM KCl, 15 mM $BaCl_2$, 5 mM EGTA, pepstatin A (0.1 μg/ml), aprotinin (1 μg/ml) and soy trypsin inhibitor (0.1 μg/ml). The suspension was mixed by inversion for 2 hrs. The resin was collected by a 5 min. slow speed (100g) centrifugation, transferred into a gravity column and washed 2 times with 10X column volume with washing buffer (WB) containing 20 mM Tris-HCl pH 8.0, 320 mM KCl, 15 mM $BaCl_2$, 5 mM EGTA, pepstatin A (0.1 μg/ml), aprotinin (1 μg/ml), soy trypsin inhibitor (0.1 μg/ml) and 0.2%DDM/0.04%CHS. On the column, cleavage of the protein was achieved by incubating the resin with WB buffer supplemented with PreScission protease to an approximate molar ratio of 20:1, followed by overnight incubation at 4°C under slow rocking motion. Cleaved protein was collected, concentrated, and ran through a Superose 6 column equilibrated in SEC buffer (20 mM Tris-HCl pH 8.0, 320 mM KCl, 15mM $BaCl_2$, 5 mM EGTA, 20 mM DTT, 2 mM TCEP, 0.025% DDM, pepstatin A (0.1 μg/ml) and aprotinin (1 μg/ml). The fractions containing tetrameric *Aplysia* Slo1 channels were concentrated to approximately 7-10 mg $ml^{-1}$ using an Amicon Ultra centrifugal filter (100-kDa MW cutoff). Within the same day of purification, samples were further used for incorporation into lipid nanodiscs.

**Reconstitution of Channel in nanodiscs**

Membrane scaffold protein MSP1E3D1 was expressed and purified from *Escherichia coli* as previously described[3], and *Aplysia* Slo1 tetramers were incorporated into lipid nanodiscs following the published protocol[4]. Lipid stock was obtained from Avanti Polar Lipids. Lipids were evaporated to remove their chloroform solvent. Dry lipid films were resuspended in water and detergent (DDM) to a final DDM concentration of 25 mM. The lipid/detergent mixture was sonicated until the solution became nearly transparent and aliquots were frozen at −80°C .

Slo1, MSP1E3D1, and lipid were mixed at a molar ratio of 1:2:85, respectively, and incubated on ice for 30 min. Detergent was removed by adding Bio-Beads SM2 to 20 mg $ml^{-1}$, followed by gentle agitation for 3 hours. The Bio-Beads were then removed by passage through a PolyPrep column, and the flow-through was centrifuged (10,000g) before size-exclusion chromatography. Ultracentrifuged samples were purified by size-exclusion chromatography on a Superose 6 Increase 10/300 GL column (GE Healthcare), equilibrated with SEC buffer (20mM Tris-HCl pH7.5, 150 mM KCl, 15mM $BaCl_2$, 5mM EGTA, 2 mM TCEP). In the low divalent ion structure, Superose 6 column was equilibrated in SEC buffer (20 mM Tris-HCl pH 8.0, 320 mM KCl, 20 mM DTT, 2 mM TCEP, 0.025% DDM, pepstatin A (0.1 μg/ml) and aprotinin (1 μg/ml). Fractions containing the nanodisc/ channel complexes were collected and concentrated using a centrifugal filter unit (100 kDa molecular weight cutoff). The samples were transferred on ice to the Cryo-EM Facility, Center for Molecular Microscopy (CMM), Center for Cancer Research (CCR), National Cancer Institute (NCI), Leidos Biomedical Research, Inc., where Cryo-EM grids were prepared within 3 hrs from the last step of purification.

**CryoEM sample preparation and imaging**

Samples were diluted to optimal concentrations ranging from 2 mg/ml to 8 mg/ml in SEC or ND buffer. Three microliters of each sample were applied to a Quantifoil R 1.2/1.3 Cu 200 mesh grid that had been glow discharged for 5 seconds using a PELCO easiGlow system (Ted Pella, Inc.) at 15 mA and 0.3 mBar. Grids were then plunged frozen in liquid ethane with a Vitrobot Mk IV (Thermo Fisher Scientific) operated at 4°C and 100% humidity, with a blot time of 2 seconds and a wait time of 5 seconds. Grids were stored in liquid nitrogen until imaging. Cryo-EM data were collected on a Titan Krios electron microscope (Thermo Fisher Scientific) operated at 300 kV and equipped with a K2 Summit direct electron detector (Gatan). Micrographs were acquired in counted mode using SerialEM[5] v3.8 with a 3×3 multi-shot setup and a nominal magnification of 29,000×, resulting in a pixel size of 0.858 Å. A total dose of 50 $e^-/Å^2$ was fractionated over 40 frames. Data quality was monitored during collection using cryoSPARC[6] Live v3.2 preprocessing.

**CryoEM data processing and reconstruction**

Motion correction, dose-fractionated weighting, and binning of collected movie data were carried out by MotionCor2[7] v1.4. The parameters of CTF were estimated by CTFFIND4[8] v4.1. EMD-8410 was used as the template for template-matching particle-picking in RELION v3.1[9]. This set of particles was classified in 2D iteratively to produce a set of 2D templates for another round of automated particle-picking. To clean up the data, extracted particles were classified in 2D for as many rounds as necessary in cryoSPARC v3.2. *Ab* *initio* volumes were determined in cryoSPARC with 5 classes, followed by heterogeneous refinement. High-quality particles were pooled. High-resolution 3D volume refinement was carried out in RELION, followed by postprocessing, CTF and aberration refinement, and finally Bayesian polishing. These "shiny" particles were transferred to cryoSPARC again for *ab initio* volume determination, homogeneous refinement, and non-uniform homogeneous refinement[10]. C4 symmetry was imposed during later refinements. Local resolution was determined by cryoSPARC.

**Structural modeling, refinement, and analysis**

Maps output by cryoSPARC were sharpened by PHENIX.auto_sharpen (PHENIX[11] v1.19). *Aplysia* Slo1 structure (PDBID:5tj6) was docked into maps using ChimeraX[12] v1.2, then refined by molecular dynamics using the ISOLDE[13] v1.2 plugin. Cycles of manual model building in COOT[14] v0.9, followed by real-space refinement in PHENIX.real_space_refine, were repeated several rounds until no noticeable improvement could be observed. The whole tetramer of Slo1, generated by a 4-fold symmetry, is used for real-space refinement, but three of the four subunits are the exact symmetric copies of the sole independent subunit. Metal ions inside the selective filter are located precisely on the 4-fold symmetry axis. MolProbity[15] v4.5 was used for model validation. Figures were prepared by ChimeraX. FSC and angular distribution plots were directly output by cryoSPARC. Local resolution figures were generated by ChimeraX. Quality report for both structures are given in Extended Data Fig. 4.

**Electrophysiological characterization of $Ba^{2+}$ dual-functional effects on BK channels**

Full-length Slo1 from *Aplysia* was subcloned into pGemHE vector[16]. Next, the plasmid was linearized with NheI before performing RNA *in vitro* T7 polymerase transcription with Ambion, Thermo Fisher Scientific kit. Finally, 50 ng of synthesized RNA was injected into each *Xenopus laevis* oocyte. Oocytes were purchased from Ecocyte Bioscience.

Excised inside-out patches were obtained using borosilicate pipettes (VWR 53432-921) with tips ranging between 0.7–1 MΩ, using recording solutions containing (in mM): pipette, 40 $KMeSO_3$, 100 N-methylglucamine–$MeSO_3$, 20 HEPES, 2 KCl, 2 $MgCl_2$, 100 μM $CaCl_2$ (pH=7.4); bath solution, 40 $KMeSO_3$, 100 N-methylglucamine–$MeSO_3$, 20 HEPES, 2 KCl, 1 EGTA, and free $BaCl_2$ concentrations varying between 1 -100 μM, previously estimated using Maxchelator[17]. Currents were recorded with an Axopatch 200B amplifier using Clampex software (Axon Instruments, Molecular Devices). Solutions containing different ion concentrations were exchanged using a fast solution exchange (BioLogic RSC-200).

**Data availability**

Coordinates and maps were deposited with PDBID:7RJT and EMD 24490, and PDBID:7RK6 and EMD 24493 respectively

**References**

**Acknowledgements**

We are indebted to Rod MacKinnon and Xiao Tao for kindly providing us the full-length Slo1 from *Aplysia californica* and the GFP nanobody constructs. SS, PM and MH were supported by the Intramural Research Program of the NIH (NINDS).

#### Author contributions

MH, RC, NdV, TG, SS and PM contributed to the initial design of the project. SS performed all molecular biological and biochemistry protocols involved in the purification of *Aplysia californica* slow1 and the incorporation into nanodiscs. PM performed the electrophysiological experiments. RC, JZ data processing and model refinement. RC, PM and MH prepared the figures. All authors contributed to discussions and preparation of the manuscript. MH and RC wrote the manuscript.

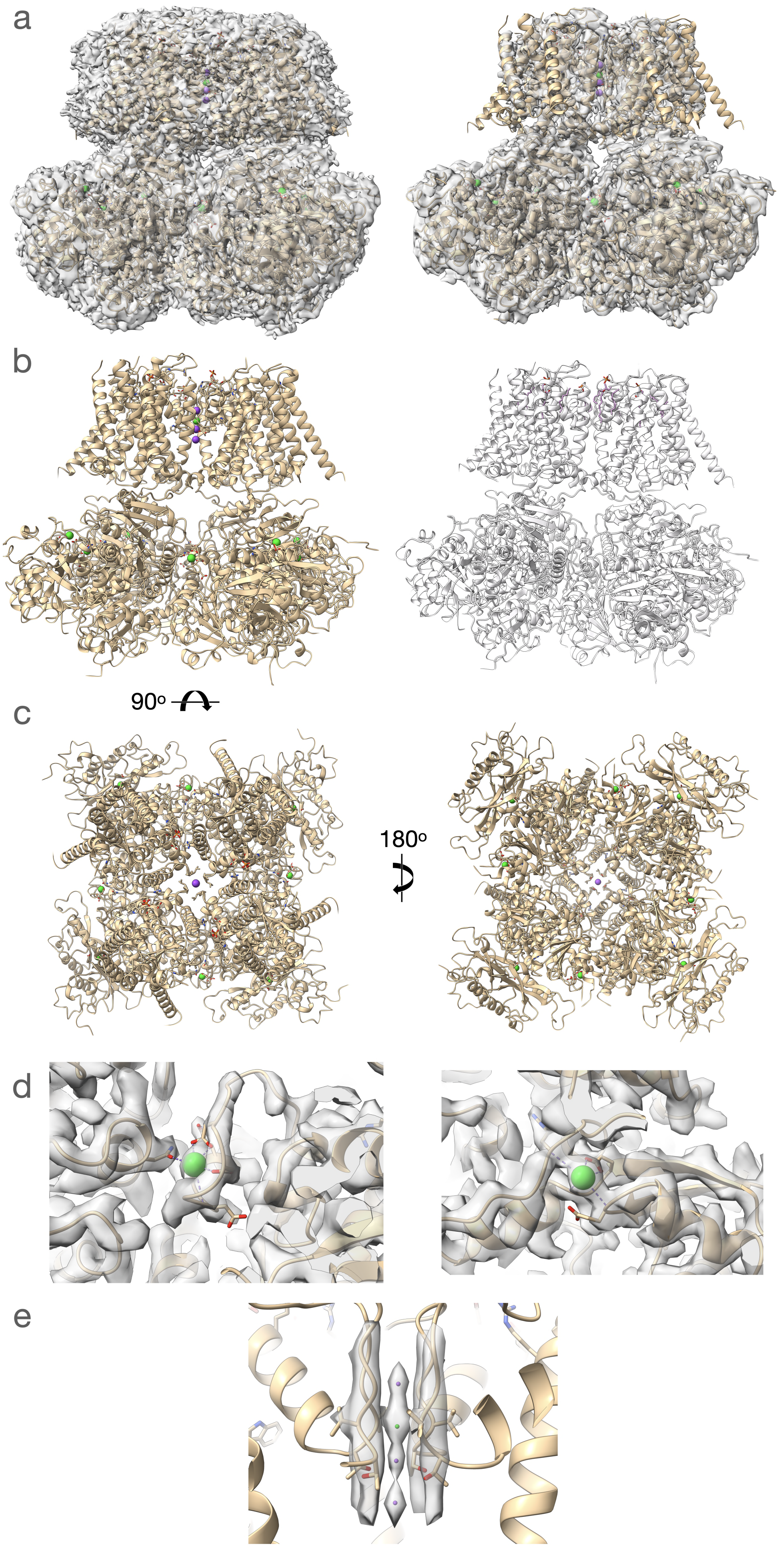

a
b
90o
c
180o
d
e

**Extended Data Fig. 1. Overview of the 10 mM $Ba^{+2}$ structure.** (**a**) overall view with map contoured to reveal the nano disc (left) and at a higher contour level (right), showing a more limited resolution of the external alpha helices in the membrane domain. (**b**) Overall view of the refined model (left). $Ba^{+2}$ ions in green. Right, a model skeleton showing the lipids identified in the density. (**c**) 90 Deg rotate views of the structure. (**d**) Detailed view of the RCK2 ($Ca^{2+}$-bowl; left) and RCK1 sites. (**e**) Selectivity filter. The map is contoured at a high counter level to reveal the $Ba^{+2}$ (green) vs. $K^{+}$ (purple) sites.

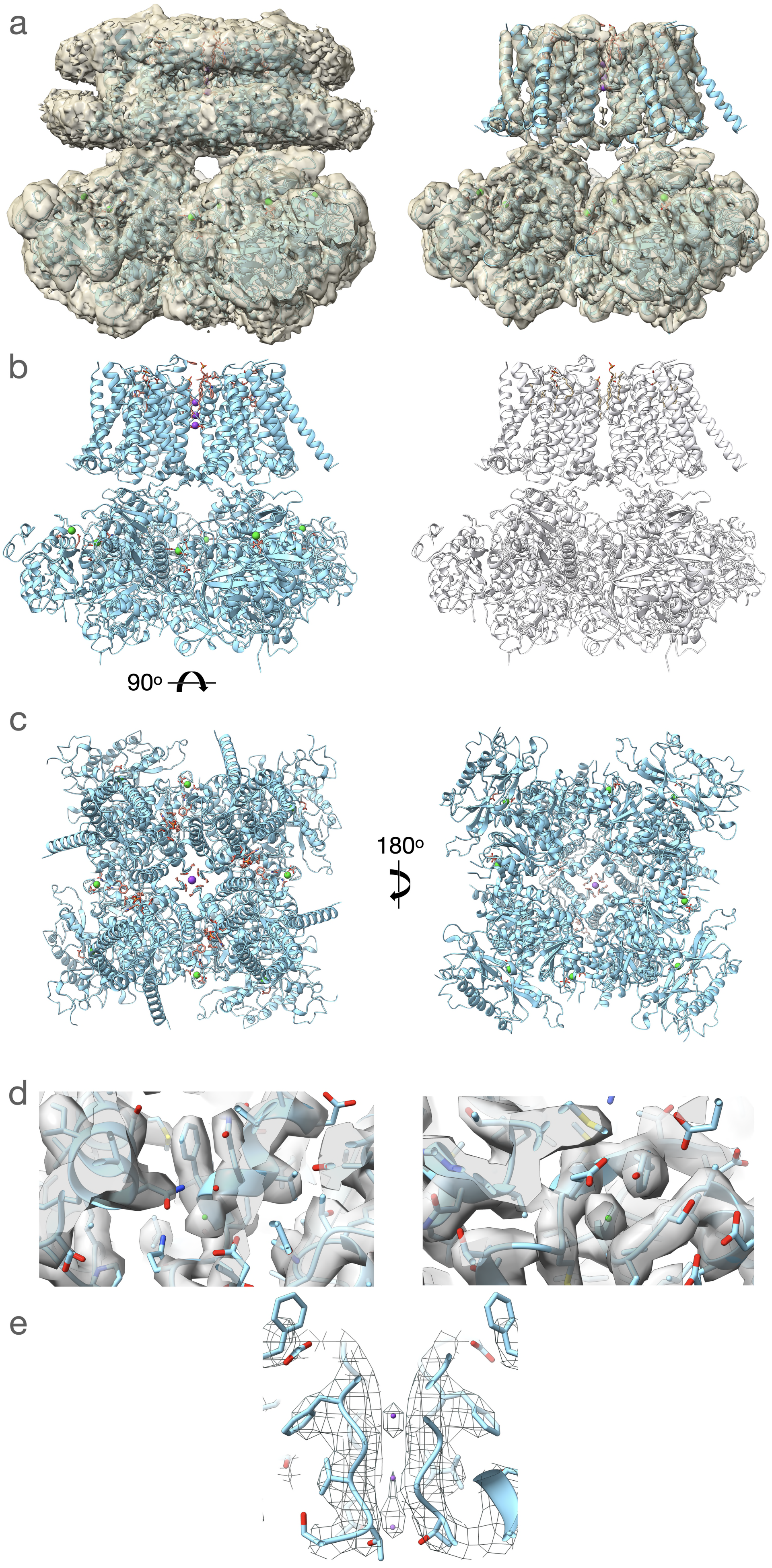

a
b
90o
c
180o
d
e

**Extended Data Fig. 2. Overview of the low-divalent-ions structure**. (**a**) overall view with map contoured to reveal the nano disc (left) and at a higher contour level (right), showing a more limited resolution of the external alpha helices in the membrane domain. (**b**) Overall view of the refined model (left). $Ba^{+2}$ ions in green. Right, a model skeleton showing the lipids identified in the density. (**c**) 90 Deg rotate views of the structure. (**d**) Detailed view of the RCK2 ($Ca^{2+}$-bowl; left) and RCK1 sites. (**e**) Selectivity filter. The map is contoured at a high counter level to reveal the $K^+$ (purple) sites.

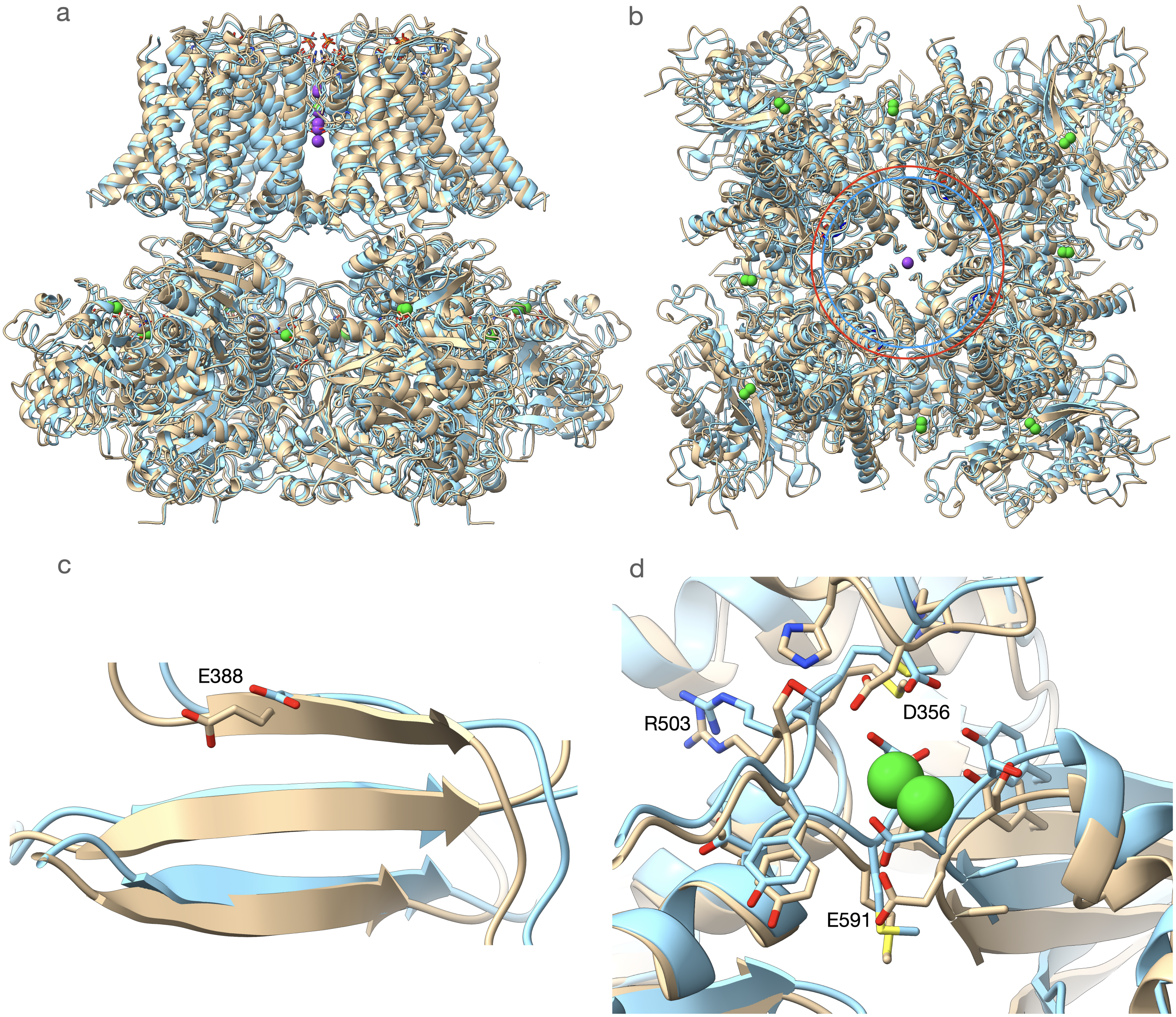
a
b
c
d
E388
R503
D356
E591

**Extended Data Fig. 3. Displacements in response to $Ba^{2+}$.** (**a**) Side view of the 10 mM $Ba^{+2}$ structure (brown) and the low-divalent-ions structure in light blue. (**b**)
Relative expansion of the intracellular domain compared using the same color scheme as in (**a**). The rings indicate Lys 320 C-alpha atom position for the 10 mM $Ba^{+2}$ structure (red ring and red atoms) and low-divalent-ions (light blue ring and blue atoms). (**c**) Zoomed in section around Glu388. The section shown is not locally aligned. For a locally aligned view see Figure 2 (main text). (**d**): Local change in the RCK1 region showing the change in the arrangement of Asp356.

1. Hite, R.K., Tao, X. & MacKinnon, R. Structural basis for gating the high-conductance $Ca^{2+}$-activated $K^+$ channel. *Nature* **541**, 52-57 (2017).
2. Tao, X., Hite, R.K. & MacKinnon, R. Cryo-EM structure of the open high-conductance $Ca^{2+}$-activated $K^+$ channel. *Nature* **541**, 46-51 (2017).
3. Inagaki, S., Ghirlando, R. & Grisshammer, R. Biophysical characterization of membrane proteins in nanodiscs. *Methods* **59**, 287-300 (2013).
4. Ritchie, T.K. et al. Chapter 11 - Reconstitution of membrane proteins in phospholipid bilayer nanodiscs. *Methods Enzymol* **464**, 211-31 (2009).
5. Mastronarde, D.N. Automated electron microscope tomography using robust prediction of specimen movements. *J Struct Biol* **152**, 36-51 (2005).
6. Punjani, A., Rubinstein, J.L., Fleet, D.J. & Brubaker, M.A. cryoSPARC: algorithms for rapid unsupervised cryo-EM structure determination. *Nat Methods* **14**, 290-296 (2017).
7. Zheng, S.Q. et al. MotionCor2: anisotropic correction of beam-induced motion for improved cryo-electron microscopy. *Nat Methods* **14**, 331-332 (2017).
8. Rohou, A. & Grigorieff, N. CTFFIND4: Fast and accurate defocus estimation from electron micrographs. *J Struct Biol* **192**, 216-21 (2015).
9. Scheres, S.H. RELION: implementation of a Bayesian approach to cryo-EM structure determination. *J Struct Biol* **180**, 519-30 (2012).
10. Punjani, A., Zhang, H. & Fleet, D.J. Non-uniform refinement: adaptive regularization improves single-particle cryo-EM reconstruction. *Nat Methods* **17**, 1214-1221 (2020).
11. Adams, P.D. et al. PHENIX: a comprehensive Python-based system for macromolecular structure solution. *Acta Crystallogr D Biol Crystallogr* **66**, 213-21 (2010).
12. Pettersen, E.F. et al. UCSF ChimeraX: Structure visualization for researchers, educators, and developers. *Protein Sci* **30**, 70-82 (2021).
13. Croll, T.I. ISOLDE: a physically realistic environment for model building into low-resolution electron-density maps. *Acta Crystallogr D Struct Biol* **74**, 519-530 (2018).
14. Emsley, P. & Cowtan, K. Coot: model-building tools for molecular graphics. *Acta Crystallogr D Biol Crystallogr* **60**, 2126-32 (2004).
15. Davis, I.W. et al. MolProbity: all-atom contacts and structure validation for proteins and nucleic acids. *Nucleic Acids Res* **35**, W375-83 (2007).
16. Liman, E.R., Tytgat, J. & Hess, P. Subunit stoichiometry of a mammalian $K^+$ channel determined by construction of multimeric cDNAs. *Neuron* **9**, 861-71 (1992).
17. Bers, D.M., Patton, C.W. & Nuccitelli, R. A practical guide to the preparation of $Ca^{2+}$ buffers. *Methods Cell Biol* **40**, 3-29 (1994).
