## Supplementary material for "Structural bases for blockade and activation of BK channels by Ba^2+^ ions": Data Quality

| | Low Ion | 10mM $Ba^{+2}$ |
|---|---|---|
| **Particles** | 973,760 | 594,822 |
| **Resolution (Å)** | 2.91 | 2.94 |
| **Cell Dimensions** | | |
| *a,b,c* (Å) | 340.5,340.5,340.5 | 292.9,292.9,292.9 |
| α,β,χ (°) | 90,90,90 | 90,90,90 |
| **Refinement** | | |
| **No. of Residues** | 3564 | 3608 |
| **RMS Bond Length (Å)** | 0.003 | 0.002 |
| **RMS Bond Angle (°)** | 0.523 | 0.478 |
| **Space Group** | P1 | P1 |
| **Ramachandran Plot** | | |
| Favored | 95.7 | 97.4 |
| Allowed | 4.3 | 2.6 |
| Outliers | 0.0 | 0.0 |
| **Molprobity** | | |
| Clash Score | 9.6 | 5.0 |
| Rotamer Outlier (%) | 0.0 | 0.0 |
| Overall Score | 1.8 | 1.4 |
| **$CC_{mask}$** | 0.79 | 0.79 |
| **PDB** | 7RK6 | 7RJT |
| **EMD** | 24493 | 24490 |

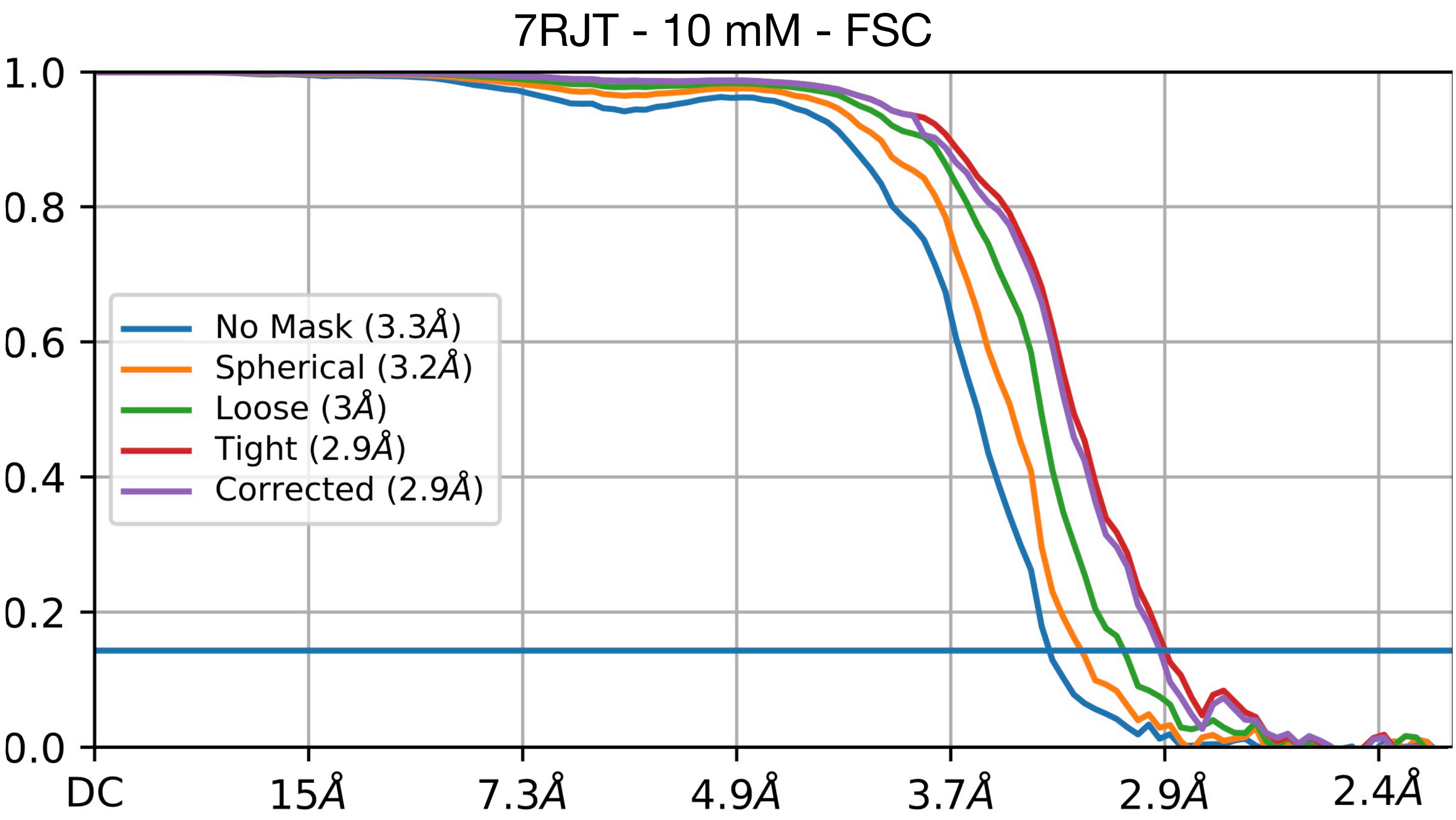

Fourier Shell Correlation Resolution 2.94A. See main text methods section.

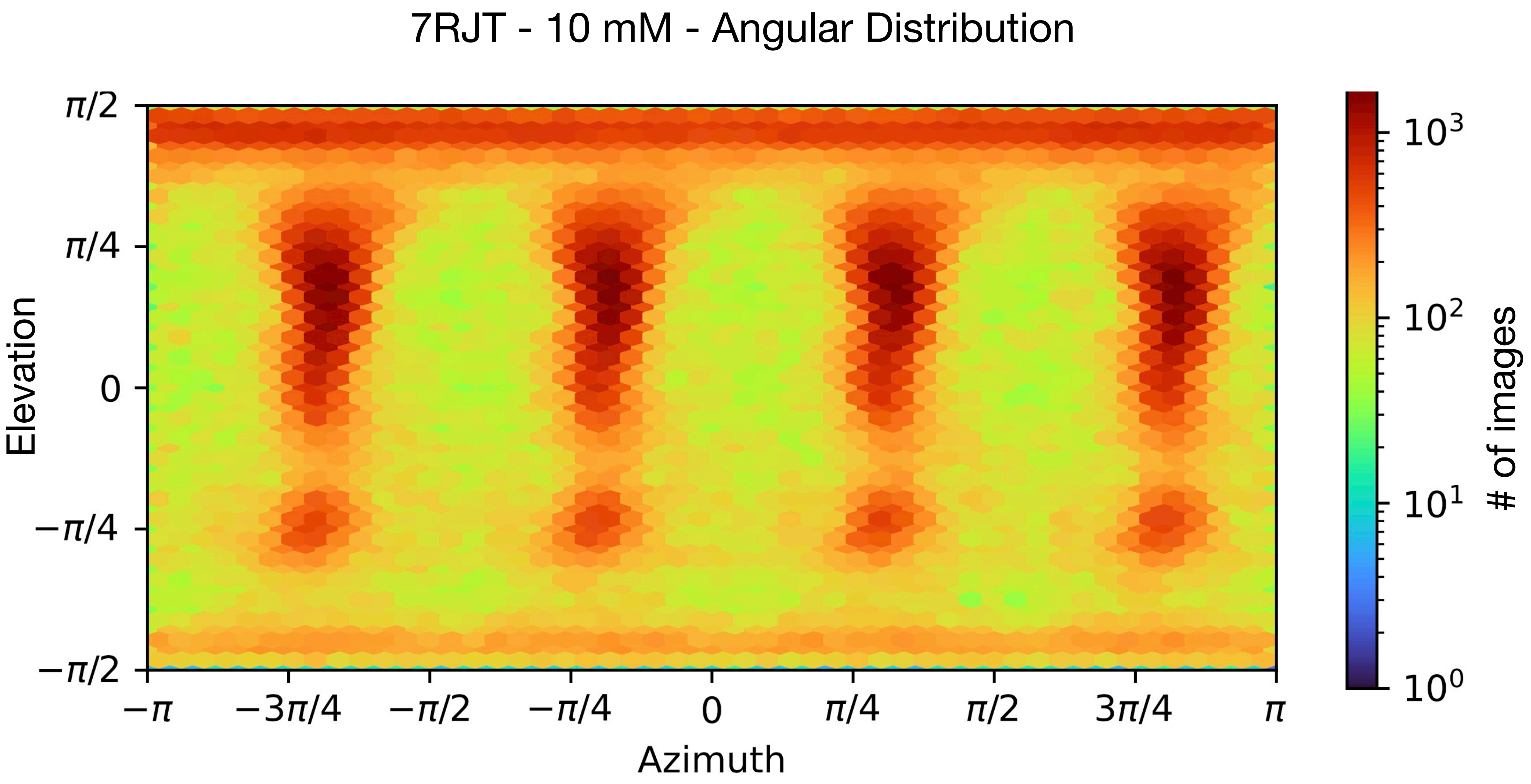

7RJT - 10 mM - Angular Distribution
π/2
π/4
0
−π/4
−π/2
Elevation
−π
−3π/4
−π/2
−π/4
0
π/4
π/2
3π/4
π
Azimuth
10³
10²
10¹
10⁰
### of images

7RJT - 10 mM - Local Resolution

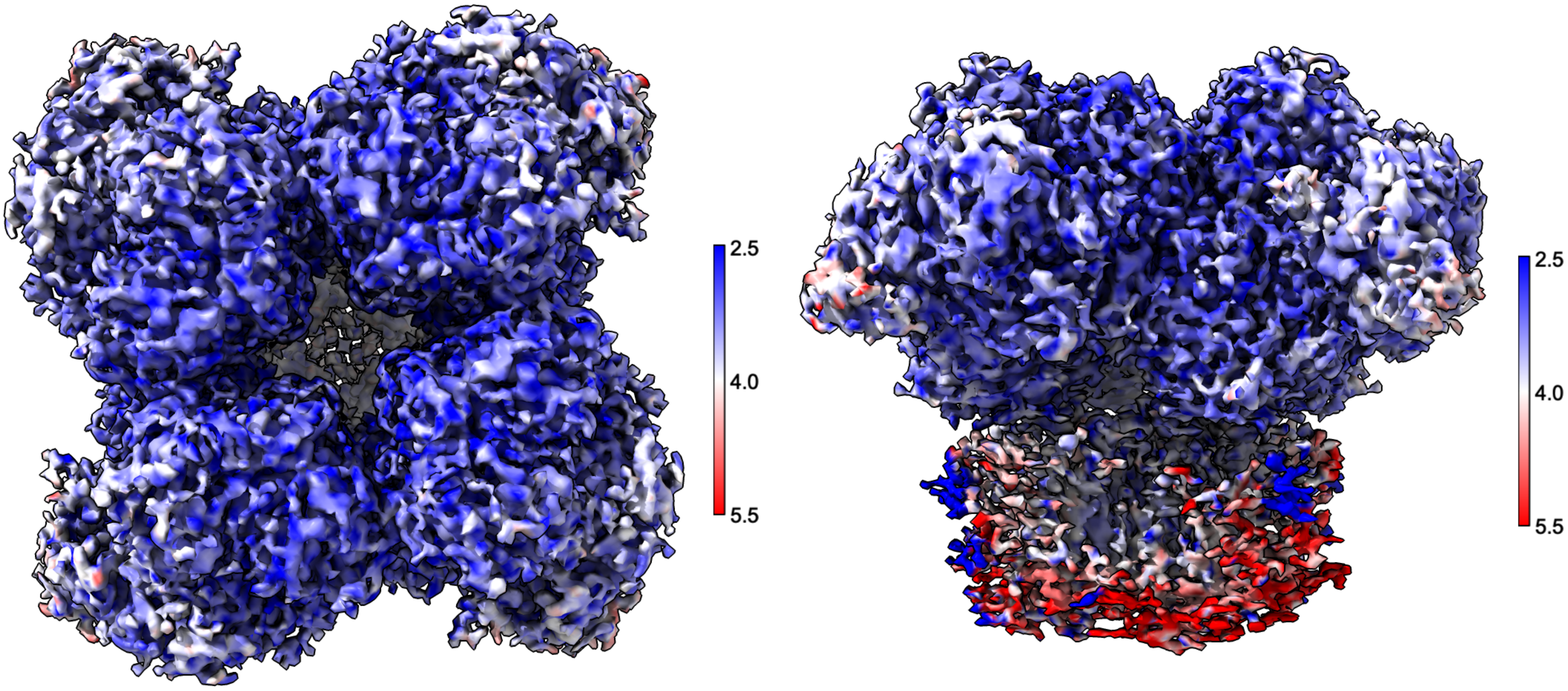

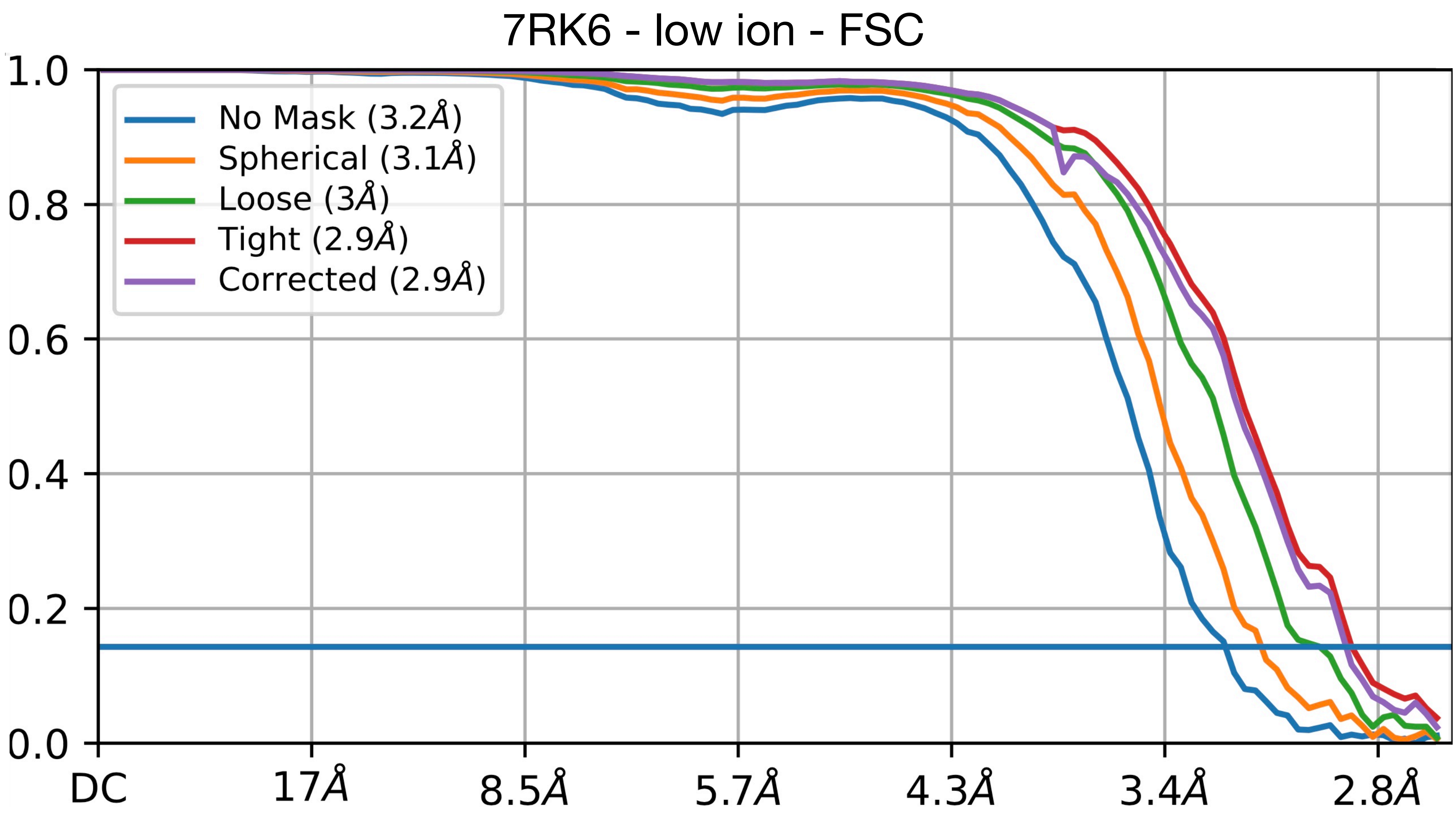

Fourier Shell Correlation Resolution 2.91A. See main text methods section.

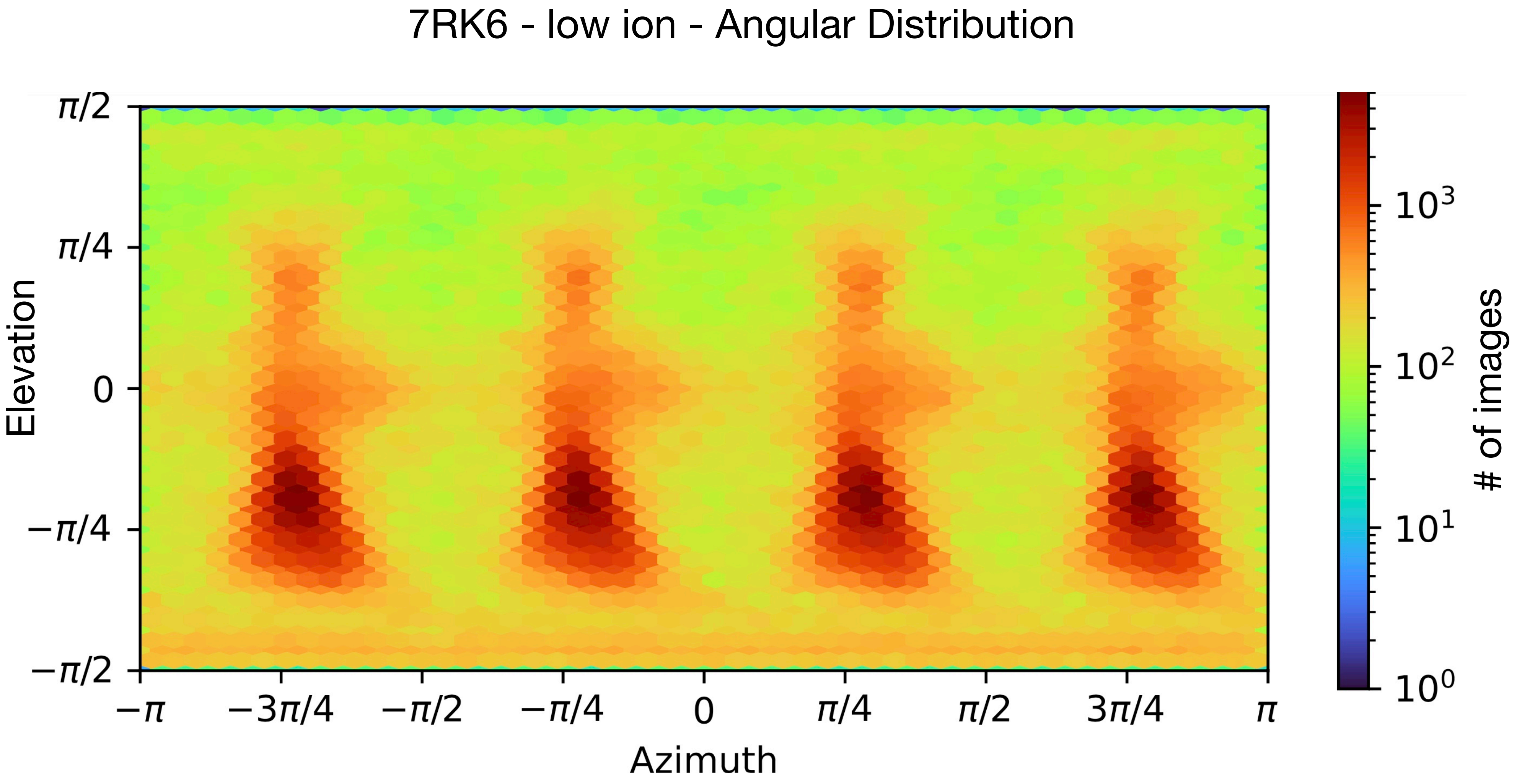
7RK6 - low ion - Angular Distribution
Elevation
π/2
π/4
0
−π/4
−π/2
−π
−3π/4
−π/2
−π/4
0
π/4
π/2
3π/4
π
Azimuth
10^3
10^2
10^1
10^0
### of images

7RK6 - low ion - Local Resolution

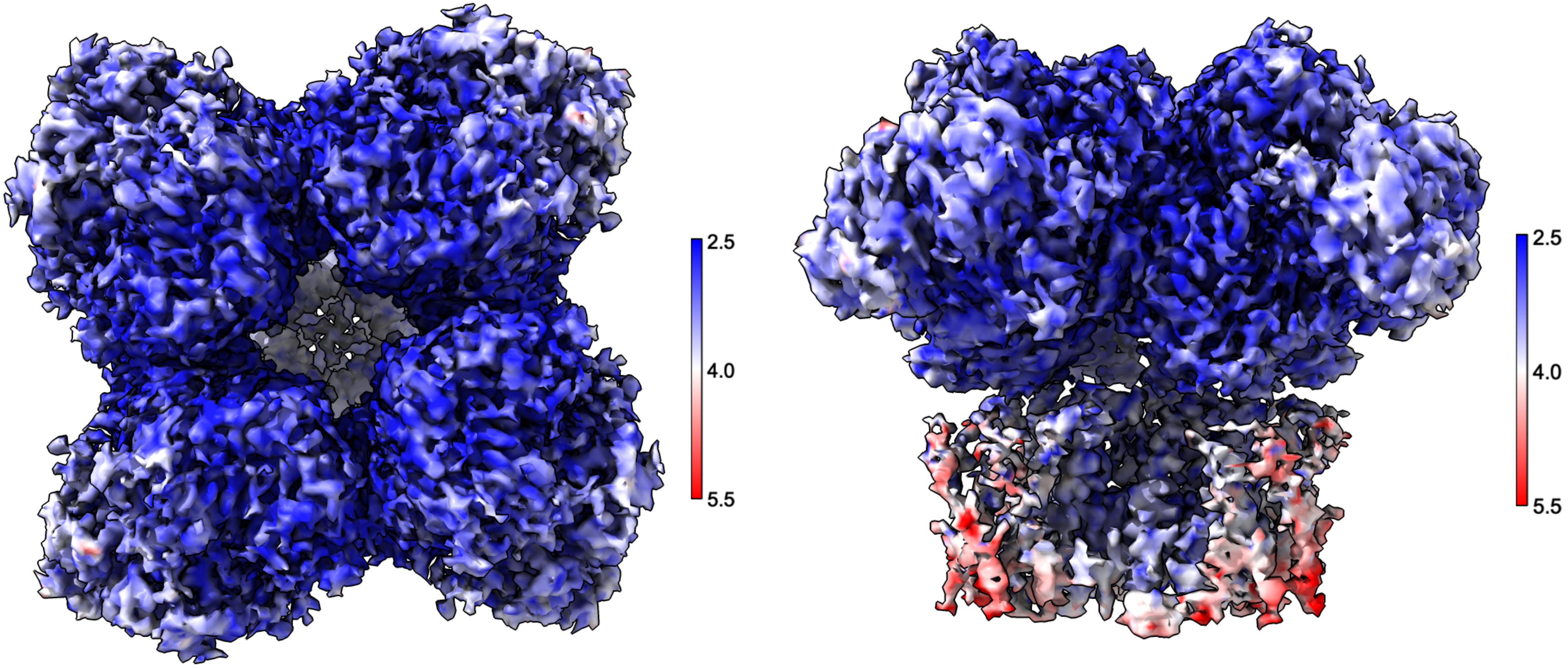
